# Automated, high-throughput quantification of EGFP-expressing neutrophils in zebrafish by machine learning and a highly-parallelized microscope

**DOI:** 10.1101/2023.08.16.553550

**Authors:** John Efromson, Giuliano Ferrero, Aurélien Bègue, Thomas Jedidiah Jenks Doman, Clay Dugo, Andi Barker, Veton Saliu, Paul Reamey, Kanghyun Kim, Mark Harfouche, Jeffrey A. Yoder

## Abstract

Normal development of the immune system is essential for overall health and disease resistance. Bony fish, such as the zebrafish (*Danio rerio*), possess all the major immune cell lineages as mammals and can be employed to model human host response to immune challenge. Zebrafish neutrophils, for example, are present in the transparent larvae as early as 48 hours post fertilization and have been examined in numerous infection and immunotoxicology reports. One significant advantage of the zebrafish model is the ability to affordably generate high numbers of individual larvae that can be arrayed in multi-well plates for high throughput genetic and chemical exposure screens. However, traditional workflows for imaging individual larvae have been limited to low-throughput studies using traditional microscopes and manual analyses. Using a newly developed, parallelized microscope, the Multi-Camera Array Microscope (MCAM™), we have optimized a rapid, high-resolution algorithmic method to count fluorescently labeled cells in zebrafish larvae *in vivo*. Using transgenic zebrafish larvae, in which neutrophils express EGFP, we captured 18 gigapixels of images across a full 96-well plate, in 75 seconds, and processed the resulting datastream, counting individual fluorescent neutrophils in all individual larvae in 5 minutes. This automation is facilitated by a machine learning segmentation algorithm that defines the most in-focus view of each larva in each well after which pixel intensity thresholding and blob detection are employed to locate and count fluorescent cells. We validated this method by comparing algorithmic neutrophil counts to manual counts in larvae subjected to changes in neutrophil numbers, demonstrating the utility of this approach for high-throughput genetic and chemical screens where a change in neutrophil number is an endpoint metric. Using the MCAM™ we have been able to, within minutes, acquire both enough data to create an automated algorithm and execute a biological experiment with statistical significance. Finally, we present this open-source software package which allows the user to train and evaluate a custom machine learning segmentation model and use it to localize zebrafish and analyze cell counts within the segmented region of interest. This software can be modified as needed for studies involving other zebrafish cell lineages using different transgenic reporter lines and can also be adapted for studies using other amenable model species.

## Introduction

Neutrophils are a subset of blood-borne polymorphonuclear leukocytes that act as a frontline defense against a wide range of insults [1]. Upon localized injury [2–4], or infection [5–7], neutrophils rapidly migrate to the affected area, where they eliminate pathogens and release factors that prime tissue repair [8,9]. Neutrophil deficiency (neutropenia), resulting from congenital conditions [10,11] or chemotherapeutic treatments [12,13], increase susceptibility to infections [14,15] and worsen the overall clinical picture of the patient. Notably, the exposure to different classes of environmentally relevant pollutants [16–18] has also been associated with neutropenia, urging the development of high-throughput assays to screen for chemicals that affect neutrophil counts. Zebrafish is now recognized as a mainstay vertebrate model to study innate immunity: the zebrafish hematopoietic program is highly conserved with higher vertebrates [19] and each spawn can yield hundreds of transparent embryos that, as early as 48 hours post fertilization (hpf), exhibit mature neutrophils [20]. Exploiting the available fluorescent reporter lines to label neutrophils *in vivo* [21,22], large cohorts of zebrafish embryos can easily be engaged in chemical-screening assays to evaluate neutrophil counts *in vivo* [23,24].

Previously published algorithms to automatically quantify fluorescent immune cells in zebrafish [25,26] relied on time-consuming positioning and imaging one larva at a time, which dramatically reduced the throughput and scalability of the assay. Improvements in technology have recently allowed for the automation of this process using scanning microscopes [27,28] which can increase the number of fish that can be feasibly examined, however restrictions still exist with these methods with regard to the time required to capture and process data as well as requisite orientation and pigmentation level of the fish, again limiting utility. Multi-camera microscope designs have been proposed previously to overcome the limited field of view of high resolution optics. Briefly, the use of multiple tightly packed microscopes in parallel can help parallelize imaging large areas such as contiguous cell culture plates [29,30] or in discrete areas such as well plates [30,31].

The recently developed Multi-Camera Array Microscope (MCAM™, Ramona Optics, Inc.) provides a novel imaging mechanism and data processing platform that overcomes multiple challenges in this workflow. An array of lenses, each coupled to a high-quality camera sensor, cover a flexibly large viewing area depending on the number of mini-microscopes employed. The system has been optimized for zebrafish behavioral and screening experiments [29,32] and functions at multiple spatial scales [30]. The MCAM™ configuration used here has 48 cameras with two distinct imaging modes, one using 24 cameras to peer into 24 wells of a 96-well plate at once with ∼3 μm per pixel resolution, and the second using 24 different cameras zoomed out to a more distant focal plane allowing the entire 96-well plate to be imaged at once with ∼9 μm per pixel resolution. In the zoomed in mode, the motorized optical head of the instrument moves in the X and Y dimensions to acquire four rapid images and yield a complete view of the plate in four seconds. The microscope stage holding the well plate moves along the optical (z) axis and controls focus of the specimen. Combining the movements of the optical head and stage, volumetric scans can be rapidly acquired in all 96 wells yielding high temporal resolution in addition to high spatial resolution throughout the 3-dimensional imaging space.

With improved imaging capabilities comes the need for efficient strategies to process the resulting data. Artificial intelligence and specifically machine learning for biomedical research is a field constantly growing in scope with novel use-cases emerging frequently. In the last decade the advent and reputable accuracies of ResNet [33] and AlexNet [34] found wide usage in image classification tasks driving applications of this type of algorithm to be developed across many computing platforms from traditional computer workstations [35–37] to smart-phones [38] to gaming systems [39]. Segmentation algorithms in parallel attempt to solve the problem of grouping semantically similar pixels within an image together, often distinguishing an object-of-interest from background [40]. In biomedical applications U-Net has been implemented repeatedly for segmentation due to its high accuracy and computational efficiency [41]. Once pixels within an image are grouped together further analysis can be focused on this region.

In this paper, we propose and demonstrate that it is possible to use an image processing pipeline based on U-Net and blob detection to algorithmically count fluorescent immune cells in zebrafish larvae. Manual counting is the current standard practice for quantification of these cells [20,42–45] while algorithmic quantification yields highly reproducible, objective values significantly faster than human analysis. This technology yields a relative measure of immune cell count and can distinguish phenotypes within a population. To this end, we treat zebrafish with both genetic and chemical immune attenuation techniques, quantify neutrophils, and compare distributions between populations by both manual and algorithmic counting. We present here an imaging methodology and open-source framework for image processing and analysis which we implement to digitally quantify the number of neutrophils present in zebrafish under experimental immunomodulatory conditions. Additionally, we validate functionality with mesh well plate inserts to expand utility in experiments requiring media exchange.

## Results

In order to develop higher throughput strategies for assessing the impact of exposure to xenobiotic toxicants and/or immunomodulatory drug candidates on neutrophil number, we partnered the high-throughput capabilities of the MCAM™ system to image zebrafish larvae in a 96-well format [29,30,32] with a transgenic zebrafish line expressing EGFP in neutrophils [22]. Our objective was to expand the tools available for using the zebrafish model in chemical and genetic screens by establishing an efficient high-throughput protocol for quantifying neutrophil numbers in transgenic zebrafish larvae.

The overall workflow includes multiple steps outlined in **Figure 1**. Transgenic zebrafish larvae are plated into 96-well plates with low autofluorescence and rapidly imaged using the MCAM™, the Z-axis is searched for best-focus frames, and then the fish is segmented from its background, and fluorescent cells are located and counted in the defined region of interest (ROI). Acquisition and processing of each 18 gigapixel dataset, representing the volume imaged across a 96-well plate, takes approximately 6 minutes with an Intel i7 12900K central processing unit (CPU) and NVidia A4000 graphics processor unit (GPU).

**Figure 1:**
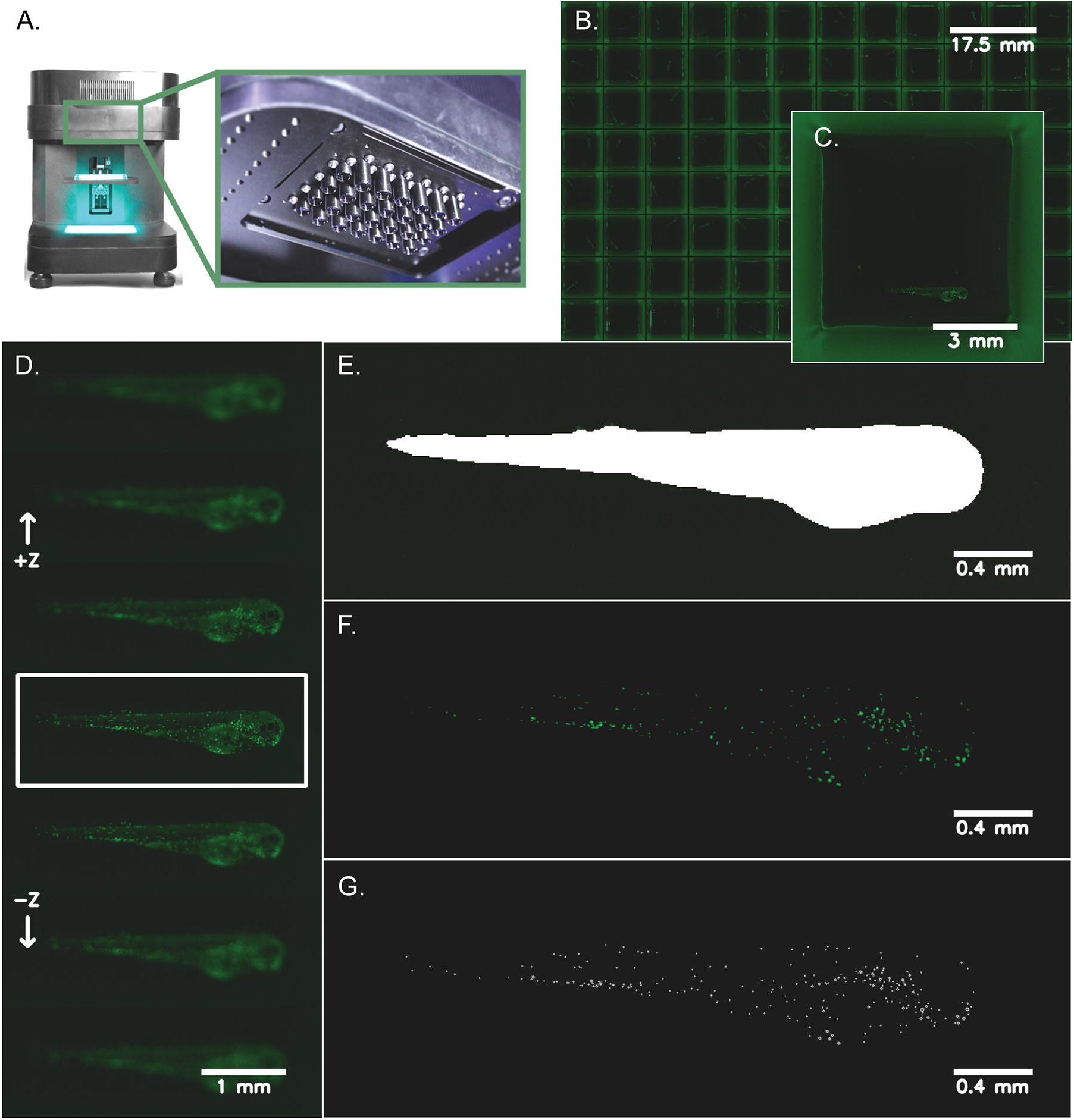
Zebrafish imaging and neutrophil quantification workflow. Transgenic zebrafish larvae (*Tg(lyz:EGFP)*) expressing neutrophil-specific EGFP were anesthetized at 72 hpf and distributed into 96-well plates with low background autofluorescence and volumetrically scanned using a MCAM™ (see **Materials and Methods**). **A**) Depicts the Multi-Camera Array Microscope (MCAM™) alongside a closeup of the 48 micro camera modules that make up the microscope array. Each lens is 12 mm in diameter. **B**) A representative image of a 96-well plate with *Tg(lyz:EGFP)* transgenic zebrafish larvae is shown. **C**) A zoomed in image (natively 3072 x 3072 x 3 pixels^2^ and ∼3 μm/pixel resolution) of a single well with a zebrafish larva in lateral orientation is shown. **D**) Following image acquisition, the Z-axis was searched automatically for the most in-focus frame of each well using a pretrained segmentation model to find a region-of-interest around each zebrafish and compute the best focus of this image region. **E)** Using the most in-focus frame for each well, each larva was segmented from the image background and a mask was generated to represent this region-of-interest. **F**) Neutrophils are shown after applying a pixel intensity threshold applied to the segmented larva which highlights the cells for counting. **G**) Individual cells were counted using blob detection techniques and are pinpointed on each image for visualization.

Machine learning based segmentation of zebrafish larvae is critical to the data acquisition pipeline as this technique is used both to define a ROI to evaluate variance of Laplacian and search the Z-axis for focus level as well as to create a ROI in which we search for cells to count. Images for training segmentation models were organized and annotated (**Figure 2A and B**) and then in order to reduce computation in this intensive step, images were downsampled for model inference, optimally on a GPU. A visualization of segmentation masks generated at different resolutions is shown in **Figure 2C**. While this strategy reduces data processing demands, and thus runtime (**Figure 2D**), it also reduces segmentation accuracy (**Figure 2E**). In developing this pipeline we found that less segmentation accuracy was needed in the first segmentation step to find best-focus frames but then higher accuracy was desired in the second segmentation step to ensure that the generated mask encompasses the entire fish and we identify all neutrophils within the region. Images were resized from 3072 x 3072 x 3 pixels to 256 x 256 x 3 for focus finding and to 1024 x 1024 x 3 for cell counting.

**Figure 2:**
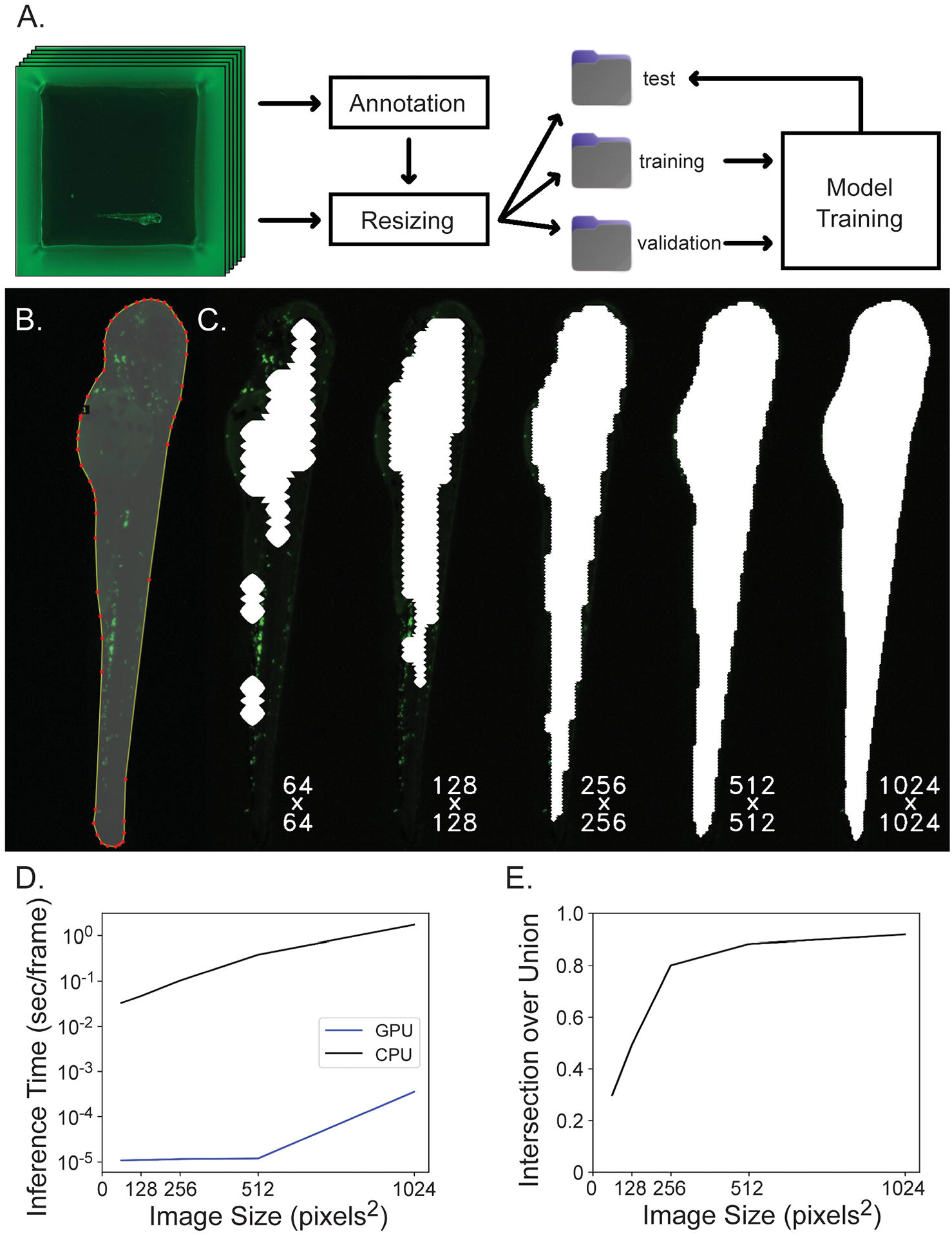
Segmentation network training and evaluation. **A**) Data is organized for model training by annotating images, resizing images and corresponding annotations to model input dimensions, separating images randomly into training, validation and test subsets and then the training and validation subsets are used for model training while the test subset is used for model evaluation. **B**) Images are annotated by outlining the fish and many of these image-label pairs are fed to U-Net to train the neural network. **C**) Square 3072 x 3072 well images are downsampled to either 64 x 64, 128 x 128, 256 x 256, 512 x 512, or 1024 x 1024 pixels^2^ to reduce computation for segmentation inference and the resulting region of interest (ROI) mask is upsampled back to the original image shape which greatly affects segmentation accuracy. Here, segmentation masks computed at different resolutions are overlaid on the original image at native resolution and cropped to display only the fish. Labels reflect the resolution downsampled to during inference. **D**) Inference time per frame is plotted against image size. Inference time increases when segmenting increasing image sizes, and this computation is completed much more efficiently on a GPU rather than CPU. The Y-axis is displayed on a log scale. **E**) Intersection over union is plotted against image size. Intersection over union improves when images are inferred at higher resolution.

Finding the optimal Z-plane and thus best focus images across 96 wells took 25 minutes for a human to manually sort through the images and select the best images from the 31 Z-planes. Segmenting fish and using the Laplacian variance algorithm within the regions of interest as described takes 3.5 minutes to choose the best frames. We compared the frames selected by algorithm versus those manually selected by a human (**Supplemental Figure S1**). 54% of the selections matched exactly and 92% of algorithm selections either matched or were within one frame of the manual selection.

Next EGFP^+^ neutrophils were counted both manually and algorithmically in zebrafish that were in a lateral orientation in the best focus frames. Manual and algorithmic counts were compared and a similar distribution and mean were found between the two quantification methods (**Figure 3A**). The number of fish both out of focus and in a non-lateral view were counted and it was found that very few fish were in these categories (**Figure 3B**). While it is not possible to count fluorescent cells in these two categories of fish we chose to disregard this factor because they made up such a small portion of the total population.

**Figure 3:**
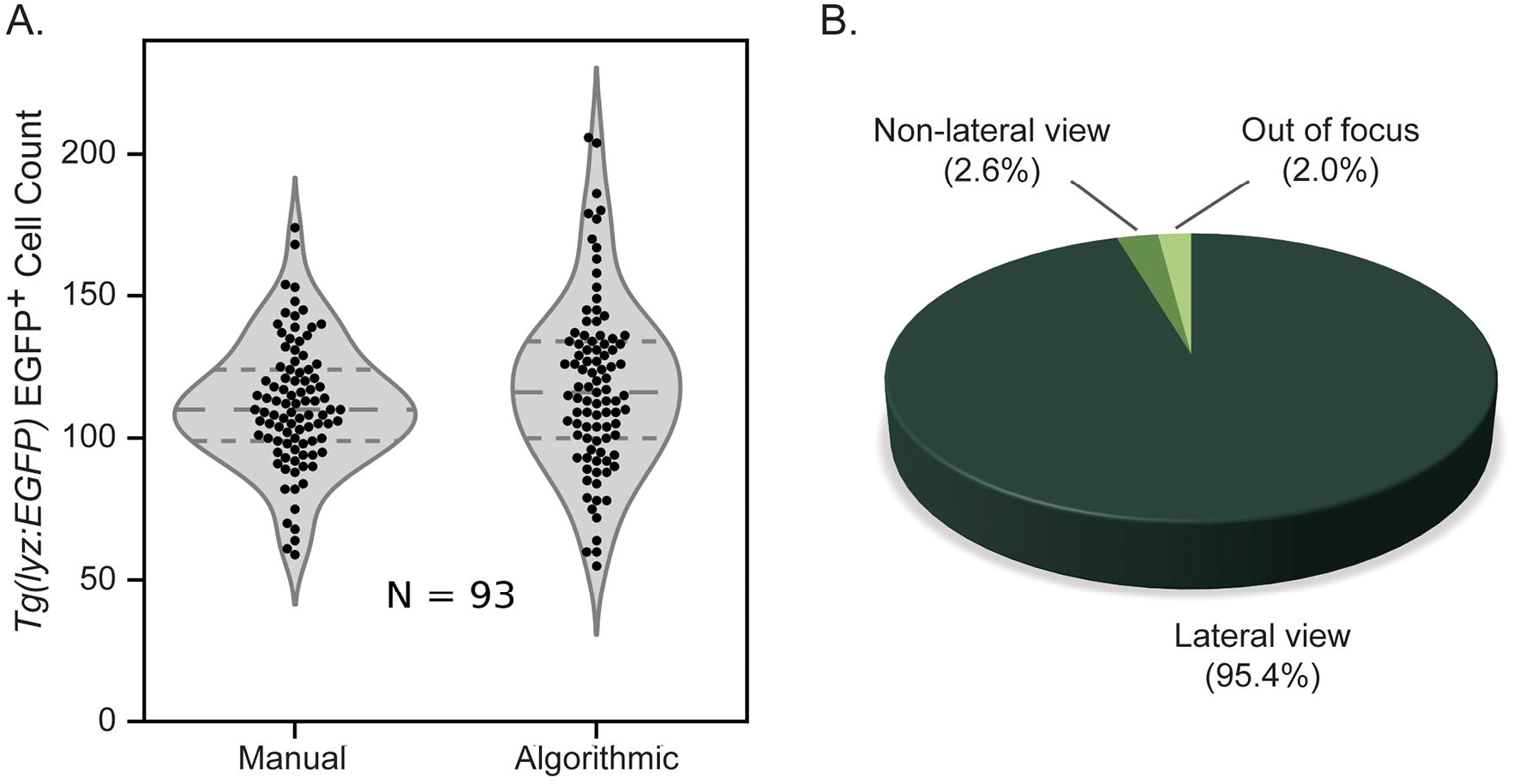
Algorithmic versus manual counting of EGFP^+^ neutrophils. **A**) Violin plot showing similarity between distribution of manual and algorithmic neutrophil counts in 72 hpf *Tg(lyz:EGFP)* zebrafish expressing neutrophil-specific EGFP with highly similar mean values (N=93). **B**) Orientation of anesthetized 72 hpf zebrafish (N=192) plated in square 96-well plates suggesting that the potential discrepancy introduced by counting cells in fish in the non-lateral orientation is minimal because this sub-population accounts for such a small fraction of the whole.

Finally, zebrafish were subjected to chemical and genetic immune attenuation methods known to reduce neutrophil numbers and EGFP^+^ neutrophils were counted for all populations including wild-type (non-EGFP) fish. The exposure of zebrafish larvae to dibutyl phthalate (DBP) is known to reduce neutrophil numbers [16] and the genetic knock-down of the *colony stimulating factor 3 receptor* (*csf3r*) gene in zebrafish larvae also has been shown to reduce neutrophil numbers [46]. When these methods were applied to *Tg(lyz:EGFP)* larvae, both methods reduced the number of EGFP^+^ neutrophils (**Supplemental Figure S2**) and no significant difference was found between manual and algorithmic counting methods using an independent t-test for all experimental conditions except wild-type fish (**Figure 4A**). A very small significance was found in the difference between the manual and algorithmic counts of wild-type fish because the manual count was zero for all fish while the algorithm counted 1 to 3 neutrophils for a few fish which turned out to be autofluorescent melanophores. Comparing any value to zero yields a significant difference. Additionally manual and algorithmic cell counts were compared, pooling the data from all experimental conditions and a linear regression resulted in an R^2^ value of 0.8974 suggesting a strong correlation between manual and algorithmic cell counts (**Figure 4B**). These results together confirm that the proposed algorithmic counting strategy is successfully matching the accuracy of manual counting.

**Figure 4:**
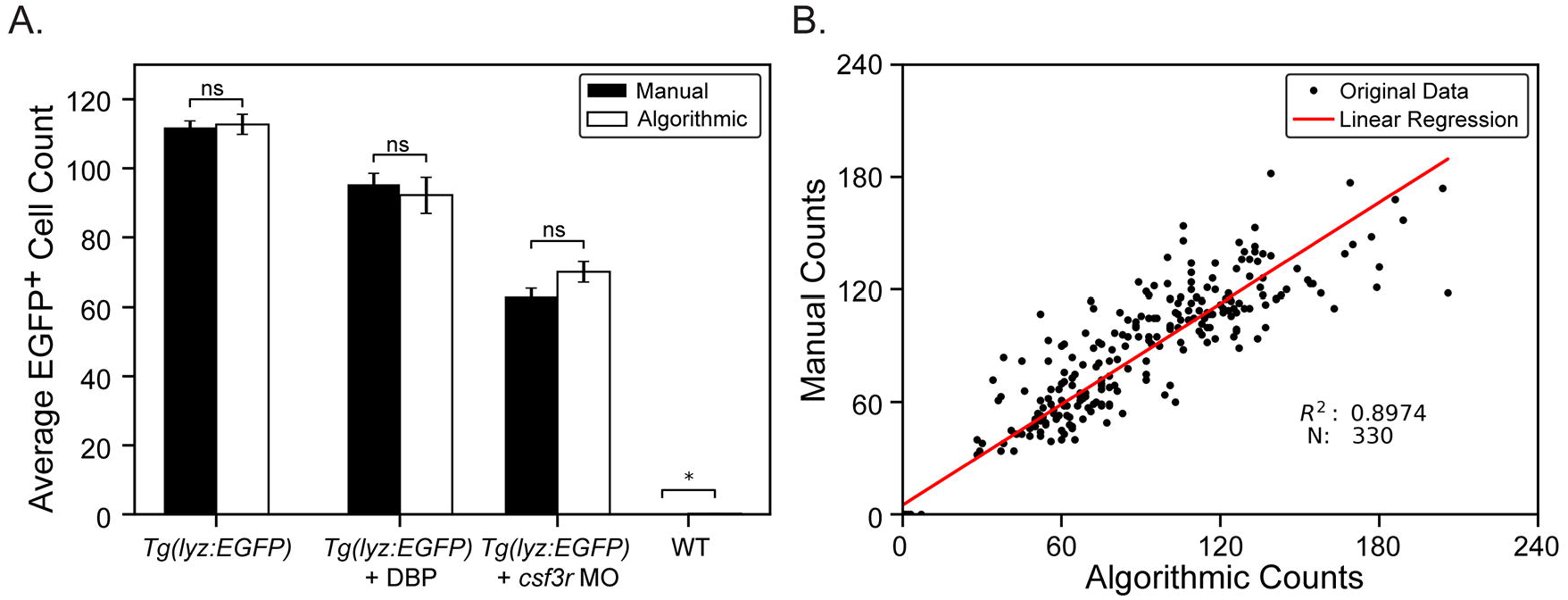
Algorithmic versus manual cell counting for experimental conditions. **A**) Knockdown and chemical modulation of zebrafish neutrophil counts. A *csf3r* antisense morpholino (MO) was injected into one-cell stage zebrafish embryos reducing neutrophil counts at 72 hpf (N = 95). Another subset of zebrafish were treated with 2 μM dibutyl phthalate (DBP), from 6 to 72 hpf, also reducing neutrophil counts but by a more subtle degree (N = 23). Neutrophil counts were obtained manually and by using the algorithmic pipeline and compared for all groups including untreated *Tg(lyz:EGFP)* (N = 93) fish and non-EGFP wild-type (WT) fish (N = 96). Data points show average neutrophil count and error bars represent the standard error of each experimental group. p-values were computed using an independent t-test. * = p ≤ 0.05, ns = no significance. **B**) Linear regression displaying strong correlation between manual and algorithmic counts for all conditions.

Once this workflow was established the imaging protocol was repeated using zebrafish in mesh well inserts to determine if they would be compatible with cell counting. Mesh well inserts have proven useful for chemical screens as they facilitate medium exchange and wash procedures [47,48]. In these mesh well liners we found many more fish (31.7% vs. 2.6% in the square well plates) to be in an orientation unsuitable for cell quantification (**Supplemental Figure S3A**) either because they were non-lateral or did not lie flat and thus had multiple best focal planes, however once these fish were removed from the dataset a strong correlation was again found between manual and algorithmic counts (**Supplemental Figure S3B**). Note that the pixel intensity threshold was reduced to 30 from 55 for these studies because the mesh, visible through the fish, altered the distribution of pixel intensities within the fish ROI. The *csf3r* morpholino injection was repeated and again neutrophil reduction was quantified by algorithmic and manual counting with similar results to the square well plate experiments (**Supplemental Figure S3C**) suggesting that mesh well inserts are amenable to this quantification workflow however there is an additional step requiring human intervention to exclude fish in non-optimal orientations.

## Discussion

Through this work we have demonstrated an automated, high-throughput neutrophil counting protocol in zebrafish larvae that can be readily adapted for use with other cell types and other model organisms. Using the MCAM™ we employ volumetric scans over an entire 96-well plate to ensure that we capture at least one in-focus image of each zebrafish. Custom neural network segmentation models yield highly accurate localization of fish and cells are counted using traditional image processing techniques and a blob detection strategy. It should not be expected that a human and computer will interpret pixel intensities and blob boundaries identically and so it is unlikely that manual and algorithmic counts will ever match up perfectly, however, using the statistical advantage we have in parallelizing many zebrafish experiments at once we yield a relative measure of neutrophil count within a population that is proven here useful in identifying differing immune cell counts between control and immune attenuated fish. Through this approach we found strong correlations between average algorithmic and manual cell counts (**Figure 4A**) with no statistically significant difference for each condition except for wild-type (non-EGFP) fish counts because the algorithm counted one to three cells in a few fish (autofluorescent melanophores) and when comparing to the manual count of zero, any small difference is found to be significant. Furthermore, we found a strong correlation of absolute cell counts between fish from all experimental conditions resulting in a linear regression R^2^ value of 0.8974 (**Figure 4B**).

Volumetric acquisitions used in conjunction with convolutional neural networks are well described in image reconstruction, composite, and focal stacking and searching techniques [37,49–51] and have proven useful for automated disease diagnosis [52,53]. By utilizing the rapid imaging of the MCAM™ throughout this 3-dimensional imaging space we are able to ensure that we capture an in-focus image of nearly every fish and minimize the temporal difference between imaging the first and last fish which is important when considering rapidly developing larvae. Once images were acquired, we implemented an automated approach for selecting the best focus frame for each well, from the Z-stack and we chose to measure focus level of an image by maximizing the variance of the Laplacian transform of the image [54], or in our case, of a region of an image. For each well, we use a custom segmentation algorithm to locate the zebrafish in each frame of its Z-stack and define a region of interest (ROI) in which the variance of the Laplacian is calculated, selecting the maximum as our best-focus frame. By considering only the variance of Laplacian of the given ROI, we can hone in on the specific focal plane of interest to ensure the zebrafish is in focus for further detailed analysis. Very few fish images (2.6%, N=192) were found to be in a non-lateral orientation unsuitable for neutrophil quantification (**Figure 3B**) and so we chose to disregard this factor in our analysis. Both the segmentation algorithm (**Figure 2C, 2E**) and the focus selection functionality (**Supplemental Figure 1**) were shown to be quite accurate.

In constructing this data processing pipeline two parameters, the pixel intensity threshold and blob size, required special consideration. When thresholding pixel intensity values we chose to set the threshold at 55 because within the fish ROI, only neutrophils have intensities above this value. We also chose to use a thresholding method which sets pixel values below the threshold to zero while values above the threshold are unmodified as opposed to binary methods which set values above the threshold to 255, assuming an 8-bit scale, which would disrupt the cell boundary that is less saturated than the centroid of each cell. The blob counting algorithm uses a blob size parameter to define the size of a gaussian kernel which is convolved with each image to find representative blobs. Using the size of neutrophils in our images we were able to set the blob size parameter at 0.05 so that individual cells are located but clusters are not found to be individual cells. Both the pixel intensity threshold and blob size values are dependent on imaging and lighting conditions and thus require constant imaging parameters to compare results between acquisitions. For example, by modifying our protocol to use plates with mesh well inserts, the pixel intensity threshold needed to be reduced because the background mesh is visible through the fish (**Supplemental Figure S4**) which alters the pixel intensity distribution. By changing the blob size parameter, other cell types with different dimensions could be quantified in the future.

A frequent question in digital cell counting is how a protocol rationalizes occluded cells or groups of cells with indistinguishable borders. Implementations can be found where researchers have used more advanced computational methods and counting strategies such as you-only-look-once (YOLO) convolutional networks [25,26,55], outlier rejection based on cell size or fluorescence intensity and watershed segmentation to define cell borders [56–58] which can yield improvements and high accuracy in terms of resolving individual cells. In our case, making use of the high parallelization of our imaging, we only require an estimation of cell count over a population and not an absolute value on a per fish basis and so further counting methods have not been found necessary.

A similar experimental protocol for fluorescent cell counting was recently devised using the WiScan Hermes High Content Imaging System and accompanying software with the goal of producing high throughput assays [28]. In this example the group reports that it takes 15 minutes to image one 96-well plate, 20 minutes to process and 10 minutes to analyze the resulting data. This method requires brightfield imaging prior to fluorescence exposure for proper segmentation of the fish and Z-stacks were again employed to acquire 5 focal planes through an overall imaging height of ∼250 micrometers. Prior to imaging, fish are dechorionated, treated with phenylthiourea to inhibit pigment formation, loaded in an alignment plate to ensure fish orientation and centrifuged, and before analysis the user inputs maximum and minimum size thresholds to assist the segmentation algorithm in finding anatomical regions of the fish. In contrast, our imaging encompasses 3.1 mm along the Z-axis, imaging in 75 seconds, a 6x increase in imaging space in 12x less time, resulting in an effective speed-up to imaging of approximately 72x across a 96-well plate. Imaging all fish quickly is critical to ensure the temporal uniformity of an experimental time point in studies focused on any level of embryonic development. Image acquisition, and the processing and analysis pipeline, designed here, have a combined runtime of 6.25 minutes compared to 45 minutes for the previous method. It is worth mentioning that we imaged twice the height that we needed to to capture an in-focus frame of each well (**Supplemental Figure 1**) and so for an optimized workflow imaging 1.5 mm instead of 3.1 mm we could reduce the full acquisition and processing time from just over six minutes, to three. When comparing algorithmic cell counts to manual we report a strong correlation across the whole fish as opposed to solely the tail region and require no user inputs. The protocol proposed here does not require any special rearing procedures and only minimal preparation ensuring that fish remain in conditions as close to natural as possible while maintaining a high-throughput workflow. We present this work as an open-source platform that researchers can adapt to their methods at no cost.

Building software and scientific technologies such as this automated neutrophil counting methodology relies on the contributions of numerous other open source softwares such as Python and the many resulting libraries as well as individuals who contribute online examples and forum assistance. With these scripts on GitLab, a user can train a custom segmentation algorithm to find a fish or any object of interest in images. An example is given for automated evaluation of the trained neural network and images can be loaded from either a pre-extracted folder or from an N-dimensional array data structure previously loaded into memory. A method for searching the Z-dimension of a volumetric stack and computation to find the best focus within a region are implemented. Finally, a few conventional image processing and quantification techniques are used to identify and count the EGFP^+^ cells, which more broadly can be used to isolate and count blobs of any identity in an image. By combining these techniques, and utilizing the MCAM™ for imaging, we demonstrate a rapid, high-throughput, cell counting process that can be easily adapted for other applications maintaining unbiased, untiring, statistical significance.

## Materials and Methods

### Zebrafish husbandry

Zebrafish husbandry and all experiments involving live animals were approved by the North Carolina State University Institutional Animal Care and Use Committee. *Tg(lyz:EGFP)*^nz117tg^ [21,22] adult zebrafish were maintained in a recirculating aquarium facility (Aquatic Habitats, Apopka, FL, USA) at 28 °C with a 14[hr light/10[hr dark cycle and fed a commercial grade zebrafish diet. Wild-type zebrafish were originally purchased from LiveAquaria (Dayton, OH, USA) and Doctors Foster and Smith (Rhinelander, WI, USA) and maintained and bred in-house for >5 years. Zebrafish embryos were obtained by natural spawning [59]. Embryos were transferred to and maintained in 100 mm Petri dishes in 1x E3 medium [60] in ultrapure water prior to imaging.

### Morpholino injection and dibutyl phthalate exposure

A *csf3r* morpholino, (5′-GAAGCACAAGCGAGACGGATGCCAT-3′) (GeneTools LLC, Philomath, OR, USA) was injected in the yolk of 1-cell stage *Tg(lyz:EGFP)* embryos as previously described [46]. Dibutyl phthalate (DBP, #36736, Millipore-Sigma, St. Louis, MO, USA) was pre-diluted in DMSO to a 10 mM concentration and stored at 4 °C. Embryos were exposed to 2 μM DBP in E3 medium from 6 to 72 hpf, with daily 99% medium change [16]. Control embryos were exposed to DMSO alone.

### Image acquisition

Transgenic zebrafish larvae expressing the neutrophil-specific *Tg(lyz:EGFP)* transgene were anesthetized at 72 hpf using 160 μg/mL tricaine (Millipore-Sigma, St Louis, MO, USA), plated into square well 96-well plates (Cytiva, Marlborough, MA, USA) and imaged using a Multi-Camera Array Microscope (MCAM™, Ramona Optics Inc., Durham, NC, USA) [29,30,32]. The MCAM™ was configured such that 24 camera sensors each image a different well of a 96-well plate. Each of the 24 cameras was set to capture an image with a total pixel count of 9.4 megapixels with an approximate resolution of 3 μm/pixel thus capturing an entire well in a single field of view. To capture the images of the other 72 wells, the MCAM™ imaging head was repositioned three times. Integrated reflection fluorescence illumination uses 450 nm LED lighting (LXML-PR02-A900, Lumileds, Netherlands) with a 495 nm short-pass excitation filter and 535/50 nm emission filters (CT495SP and ET535/50m, Chroma, Bellows Falls, VT, USA). The microscope stage, holding the well plate, was moved in the Z dimension and the optimal focal plane was determined by eye. From this plane the Z-stage was moved up and down to determine travel range extrema where all fish would be out of focus and it was found that 15 Z-slices (100 micrometer axial step per slice) on either side of the optimal focal plane would guarantee that the MCAM™ captures an in-focus image of the fish. Four axial (z) stack acquisitions were quickly captured, one at each of four lateral (x, y) locations to image the full well plate at all relevant heights in 75 seconds. Thirty-one Z-slices were obtained for each camera for each of the four axial stacks imaging 128 x 85 x 3.1 mm overall. Two hundred ms exposure, 50% brightness, 2.4 digital gain and 1.0 analog gain were used for all acquisitions. The four Z-stacks were automatically combined by the MCAM™ software yielding one large volumetric scan containing the 31 axial slices of all 96 wells. Data was stored in an HDF5 file containing both the raw image data, as well as the metadata that describes the imaging settings previously mentioned ensuring accurate offline analysis. Each individual well image at each slice has dimensions 3072 x 3072 x 3 pixels resulting in composite Z-slice images of 36,864 x 24,576 x 3 pixels or 906 megapixels per slice.

### Segmentation model training and evaluation

Sixty in-focus images of individual zebrafish larvae in wells were selected at random for label annotation (**Figure 2B**) and randomly sorted into segmentation model training, validation and test subsets (30, 13, and 17 images respectively) (**Figure 2A**). Custom segmentation models were trained to detect and segment fish from their background at different resolutions (**Figure 2C**) and thus locate them within their well. The overall dataset of sixty frames was divided into training, validation, and test subsets in order to properly train and evaluate the model. Individual images at 3072 x 3072 x 3 pixels were labeled by outlining the ROI (the zebrafish) using the VGG Imaging Annotator [61].

Images were downsampled to 64 x 64, 128 x 128, 256 x 256, 512 x 512, or 1024 x 1024 pixels^2^ and five different U-Net neural networks [41] were trained, one for each resolution, and each model learns its parameter weights through an iterative training process. Models were trained for fifty epochs using Dice loss [62] as the loss function, Adam [63] as the optimizer and learning rate beginning at 5E-4 and decreasing by a factor of ten both at the fifteenth and fortieth epochs. Once a model has been trained, new images that it has not seen before can be analyzed by the network resulting in statistical predictions as to where the boundary of a fish is likely to be. An image mask is generated from this information, hiding the image background as determined by the model, and thus highlighting the fish in the frame for further analysis. To evaluate a segmentation model the test dataset (N=17) was input into the model, segmented, and the intersection over union (IoU) [64] of segmentation mask and truth labeled fish was evaluated. Segmentation IoU and inference speed were evaluated and compared for five models, each trained with data downsampled to the resolutions mentioned.

### Image processing and cell counting

For each frame in each well, the first stage of the neutrophil quantification algorithm attempts to segment a fish from the background at 256 x 256 pixel resolution yielding a ROI, which is then scored by computing the variance of the Laplacian of the region [54]. One best frame, maximizing variance of Laplacian, is selected from the Z-stack for each well. Fish are segmented from the best-frames and a pixel intensity threshold and difference-of-Gaussian blob detection [65] were implemented to locate the centroid of each blob in the region. For this second step segmentation is computed at 1024 x 1024 resolution for higher segmentation accuracy. The pixel intensity threshold was determined interactively using ImageJ. The identified blobs were counted for each well.

To manually accomplish these steps, best frames were selected by a human observer for 87 wells of a 96-well plate, which contained one fish each while the remainder of the wells were empty. Cells were counted for each fish in each of the best frames using the built-in cell counter ImageJ plugin which allows the user to label each pixel that they click. Orientation of fish was manually scored as either the lateral or non-lateral orientation with lateral view defined as having only one eye visible. Cells in each successive well plate dataset were manually counted using the same strategy. For algorithmic neutrophil counts, each well plate dataset was passed through the cell counting pipeline which selected the best focus frames and then identified and counted blobs within each ROI of these frames. Cell counts from empty wells were discarded.

Neutrophil counts were compared between manual and algorithmic counting methods for each experimental condition using an independent T-test implemented with SciPy’s statistics library. Differences were considered statistically significant with p-values greater than 0.05. Error was determined by calculating the standard error of each set of cell counts. Cell counts from all experimental conditions were then pooled and linear regression, implemented with SciPy’s curve fitting module, was used to visualize the correlation of manual and algorithmic counts overall.

### Mesh well inserts

Once the workflow was established, *csf3r* morpholino injections were repeated for another group of embryos and at 72 hpf the larvae were plated into mesh insert well plates (MANM10010, Millipore-Sigma). Segmentation models were trained on the mesh well data set to recognize zebrafish with the mesh background. The imaging and quantification procedure was repeated using a pixel intensity threshold of 30 instead of 55 because the distribution of pixel intensities in images had changed.

## Data Availability

All image datasets used in these experiments are located online for viewing at: https://gigazoom.ramonaoptics.com/Neutrophil_Quantification/

Raw data is available at:

- Efromson, John. (2023). Zebrafish neutrophil counting segmentation models. https://doi.org/10.5281/zenodo.8184808
- Efromson, John. (2023). Zebrafish with lyz:EGFP expressing neutrophils: Non-injected stack [Data set]. Zenodo. https://doi.org/10.5281/zenodo.8035041
- Efromson, John. (2023). Zebrafish with lyz:EGFP expressing neutrophils: csf3r_MO injected stack [Data set]. Zenodo. https://doi.org/10.5281/zenodo.8035102
- Efromson, John. (2023). Zebrafish WT: Z-stack [Data set]. Zenodo. https://doi.org/10.5281/zenodo.8035129
- Efromson, John. (2023). Zebrafish with lyz:EGFP expressing neutrophils: Dibutyl phthalate Stack [Data set]. Zenodo. https://doi.org/10.5281/zenodo.8122771
- Efromson, John. (2023). Zebrafish with lyz:EGFP expressing neutrophils: Mesh Well Inserts Z-stack 1 [Data set]. Zenodo. https://doi.org/10.5281/zenodo.8035205
- Efromson, John. (2023). Zebrafish with lyz:EGFP expressing neutrophils: Mesh Well Inserts Z-stack 2 [Data set]. Zenodo. https://doi.org/10.5281/zenodo.8035269
- Efromson, John. (2023). Zebrafish with lyz:EGFP expressing neutrophils: Mesh Well Inserts Z-stack 3 [Data set]. Zenodo. https://doi.org/10.5281/zenodo.8035300

## Software Availability

Open source neutrophil quantification software can be found online at: https://gitlab.com/ramona-applications/neutrophil-quantification

## Acknowledgements

Research reported in this publication was supported by the Office of Research Infrastructure Programs (ORIP), Office Of The Director, National Institutes of Health (NIH) and the National Institute of Environmental Health Sciences (NIEHS) of the NIH under Award Number R44-OD024879 and by the NIEHS of the NIH under Award Number P42-ES031009. The content is solely the responsibility of the authors and does not necessarily represent the official views of the National Institutes of Health.

## Contributions

J. Efromson, G. Ferrero and J.A. Yoder contributed to the conceptualization of this project. G. Ferrero maintained zebrafish and implemented biological methods. J. Efromson, G. Ferrero, and A. Barker imaged zebrafish. A. Bègue and M. Harfouche assisted with dataset acquisition. G. Ferrero manually analyzed images. J. Efromson developed software and analyzed results. A. Bègue, T.J.J. Doman, V. Saliu, P. Reamey, and K. Kim assisted with MCAM™ integration. A. Bègue, J. Efromson, G. Ferrero and J.A. Yoder interpreted the results. C. Dugo assisted with online data presentation. All authors wrote, revised and edited the manuscript.

