## Supplemental Material for "Automated, high-throughput quantification of EGFP-expressing neutrophils in zebrafish by machine learning and a highly-parallelized microscope"

---

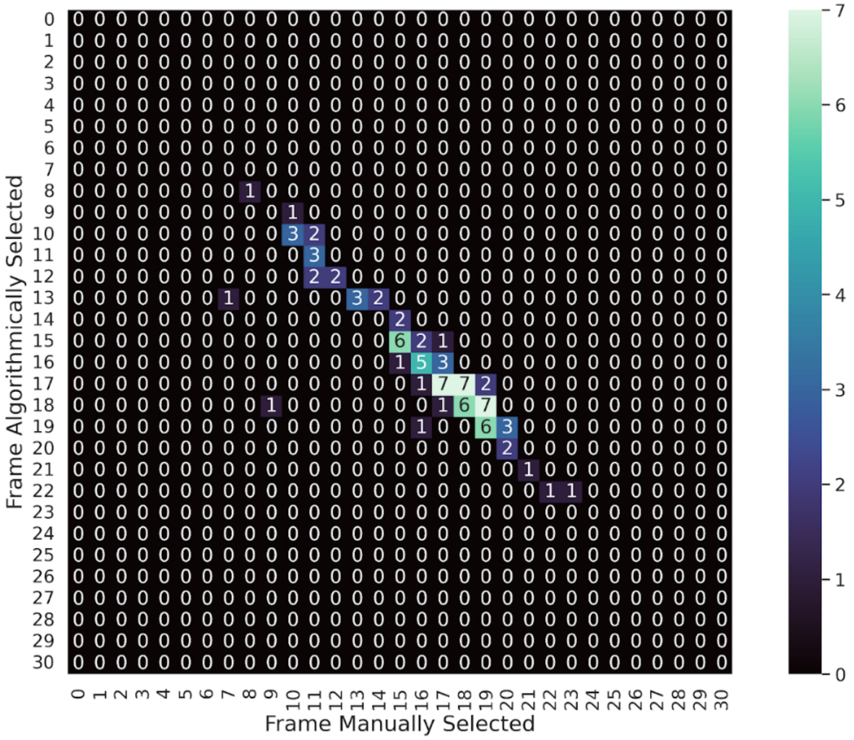

**Supplemental Figure S1: Correlation between manual and algorithmic selection of best-focus frames.**

Confusion matrix showing the correlation between manual and algorithmic selection of best-focus frames. The fish in each frame is segmented by a machine learning segmentation model and the variance of the Laplacian of this region is computed and maximized to select the best focus frame from each Z-stack. When manual selection matches algorithmic selection, counts lie along the diagonal from top left to bottom right. 54% of selections match exactly between manual and algorithmic selection while 92% of the algorithmic frame selections are within one frame of the manually selected. Data represented here is from one 96-well plate and suggests that many extra z-slices were acquired than were needed because only the center ~1.5 mm were the in-focus frames of interest.

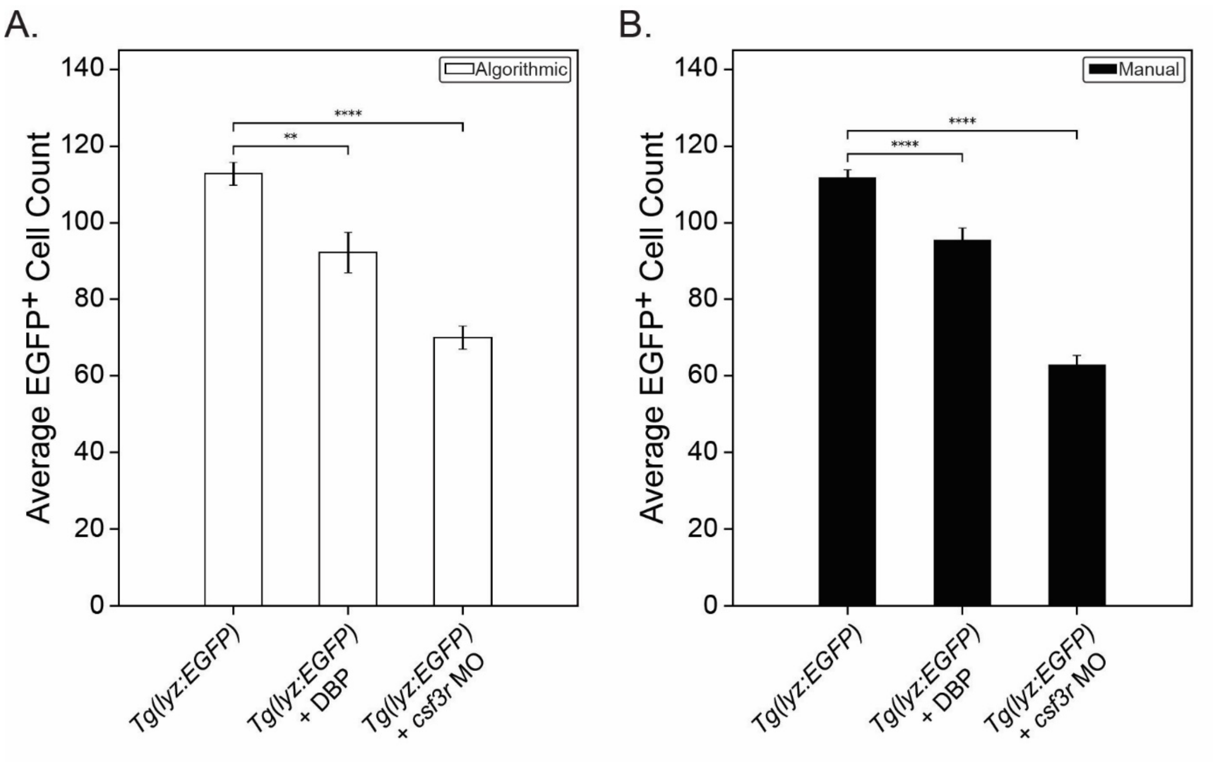

**Supplemental Figure S2: Validation of chemical and genetic methods to reduce neutrophil number.**

Knockdown and chemical modulation of zebrafish neutrophil counts. A *csf3r* antisense morpholino (MO) was injected into one-cell stage zebrafish embryos reducing neutrophil counts at 72 hpf (N = 95). Another subset of zebrafish were treated with 2 μM dibutyl phthalate (DBP), from 6 to 72 hpf, also reducing neutrophil count but by a more subtle degree (N = 23). Average neutrophil counts were compared to wild-type (WT) fish (N = 96) and the statistical significance of each method for reducing neutrophil numbers was determined using **A**) algorithmic counts and **B**) manual counts. Data points show average neutrophil count and error bars represent the standard error of each experimental group. p-values were computed using an independent t-test. ** $p \le 0.01$; **** $p \le 0.0001$.

*Note: this is the same dataset shown in Figure 4.*

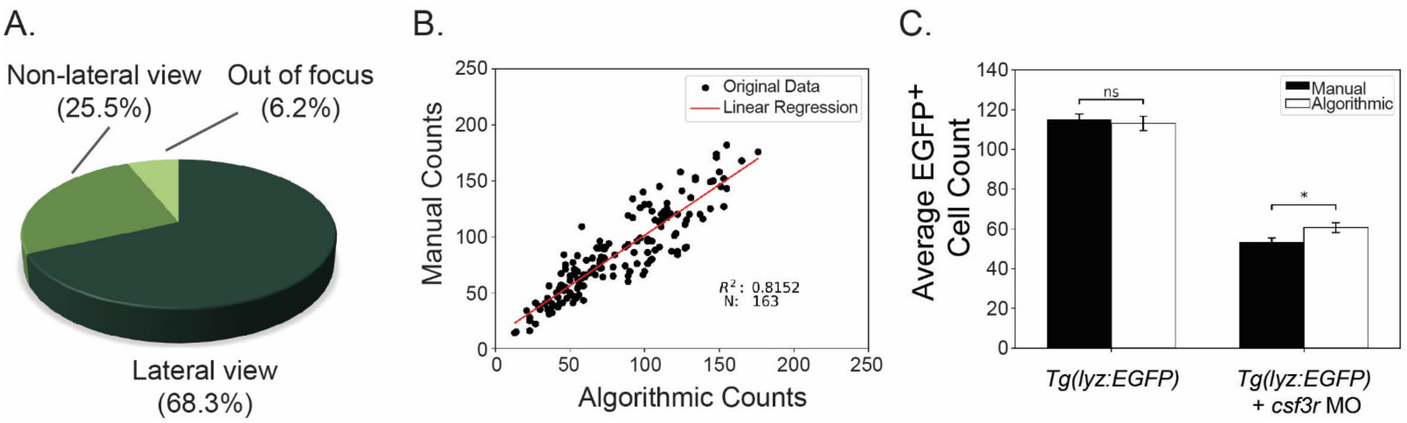

**Supplemental Figure S3: Algorithmic versus manual neutrophil counts for larvae in well plates with mesh well inserts.**

**A**) Proportion of zebrafish in the lateral or non-lateral orientation in 96-well plates with mesh inserts at 72-hpf (N=243). **B**) Linear regression displaying strong correlation between manual and algorithmic counts for *Tg(lyz:EGFP)* fish in mesh-well insert well plates (N=163). **C**) Average cell count for untreated *Tg(lyz:EGFP)* fish in mesh wells (N=76) and *Tg(lyz:EGFP)* fish injected with *csf3r* morpholino (MO) (N=87) as determined by both manual and algorithmic counting. Error bars represent the standard error of each experimental group and p-values were computed using an independent t-test. * = p ≤ 0.05, ns = no significance.

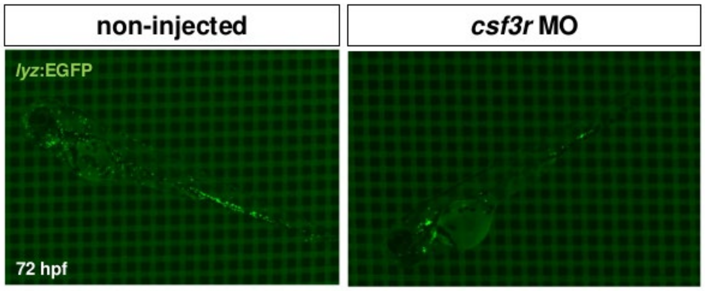

**Supplemental Figure S4: Evaluation of imaging larvae with mesh well inserts.**

Transgenic *Tg(lyz:EGFP)* zebrafish larvae (72 hpf) in mesh wells inserts in a 96-well plate. Larve were untreated (left) or injected with a *csf3r* morpholino (MO) (right). Fish exhibit a lateral orientation defined as having only one eye visible.
